# Flavivirus prM interacts with MDA5 and MAVS to inhibit RLR antiviral signaling

**DOI:** 10.1101/2022.04.25.489335

**Authors:** Liyan Sui, Yinghua Zhao, Wenfang Wang, Hongmiao Chi, Tian Tian, Ping Wu, Jinlong Zhang, Yicheng Zhao, Zheng-Kai Wei, Zhijun Hou, Guoqiang Zhou, Guoqing Wang, Zedong Wang, Quan Liu

**Affiliations:** Center of Infectious diseases and Pathogen Biology, Key Laboratory of Organ Regeneration and Transplantation of the Ministry of Education, The First Hospital of Jilin University, State Key Laboratory of Zoonotic Diseases, Changchun, China; College of Basic Medical Science, Jilin University, Changchun, China; College of Wildlife and Protected Area, Northeast Forestry University, Harbin, China; School of Life Sciences and Engineering, Foshan University, Foshan, China; The Biological safety level-3 Laboratory, Changchun Institute of Biological Products Co., Ltd., Changchun, China

**Author notes:** Correspondence: Zedong Wang or Quan Liu.

## Abstract

Vector-borne flaviviruses, including tick-borne encephalitis virus (TBEV), Zika virus (ZIKA), West Nile virus (WNV), yellow fever virus (YFV), dengue virus (DENV), and Japanese encephalitis virus (JEV), pose a growing threat to public health worldwide, and have evolved complex mechanisms to overcome host antiviral innate immunity. However, the underlying mechanisms of flavivirus structural proteins to evade host immune response remain elusive. Here we show that TBEV structural proteins, including pre-membrane (prM), envelope, and capsid proteins, could inhibit type I interferon (IFN-I) production. Mechanically, TBEV prM interacted with both MDA5 and MAVS and interfered with the formation of MDA5-MAVS complex, thereby impeded the nuclear translocation and dimerization of IRF3 to inhibit RLR antiviral signaling. ZIKA and WNV prM was also demonstrated to interact with both MDA5 and MAVS, while dengue virus serotype 2 (DENV2) and YFV prM associated only with MDA5 or MAVS to suppress IFN-I production. In contrast, JEV prM could not suppress IFN-I production. Overexpression of TBEV and ZIKA prM significantly promoted the replication of Sendai virus. Our findings reveal the immune evasion mechanisms of flavivirus prM, which may contribute to understanding flavivirus pathogenicity, therapeutic intervention and vaccine development.

**Author Summary:** All flaviviruses must overcome the antiviral innate immunity to infect vertebrate host. The non-structural proteins of flaviviruses are mainly responsible for viral replication and host innate immune escape, and the structural proteins are involved in the virus assembly, which are potential targets for prevention and treatment of flaviviral infections. Whether flavivirus structural proteins participate in host innate immune escape remains to be determined. Here, we found that tick-borne encephalitis virus structural proteins precursor membrane (prM), capsid, and envelope proteins can antagonize type I interferon production, in which prM interacts with both MDA5 and MAVS to inhibit RLR antiviral signaling. Additionally, Zika virus and West Nile virus prMs also interact with both MDA5 and MAVS, while dengue virus serotype 2 and yellow fever virus prMs associate only with MDA5 or MAVS to suppress type I interferon production. In contrast, Japanese encephalitis virus prM cannot antagonize type I interferon production. Our findings reveal that flavivirus prM inhibits type I interferon production via interacting with MDA5 and/or MAVS in a species-dependent manner, which may contribute to understanding flavivirus pathogenicity, therapeutic intervention, and vaccine development.

## Introduction

Vector-borne flaviviruses in the family *Flaviviridae* are an important source of emerging and re-emerging infectious diseases worldwide, with medically important flaviviruses of tick-borne encephalitis virus (TBEV), Japanese encephalitis virus (JEV) and West Nile virus (WNV), which cause severe encephalitis, dengue virus (DENV), and yellow fever virus (YFV), which cause hemorrhagic fever, as well as Zika virus (ZIKV), which causes severe fetal abnormalities in pregnant women and Guillain-Barré syndrome in adults [1]. In the last decades, flaviviruses have caused several epidemic outbreaks, including ZIKV, DENV and WNV [2–4]. Only for DENV, more than 40% of the world’s population is at the risk, and at least 50 million infections occur annually [5].

The type I interferon (IFN-I) is the first line of a powerful barrier against viral infections through evolutionary conserved pattern recognition receptors (PRRs), such as Toll-like receptors (TLRs), retinoic acid-inducible gene I (RIG-I)-like receptors (RLRs), NOD-like receptor (NLR), and cytoplasmic DNA receptors [6–8]. Recognition of viral RNA triggers the RLRs, such as RIG-I and melanoma differentiation–associated protein 5 (MDA5), to recruit mitochondrial antiviral signaling protein (MAVS) that stimulates the downstream TANK binding kinase 1 (TBK1) and IKKε, thereby activating the transcription factors IRF3 and NK-κB to induce interferon production [9, 10]. The secreted IFN-I binds to IFN receptor to activate Janus kinases, Jak1 and Tyk2, to phosphorylate signal transducer and activator of transcription (STAT)1 and STAT2, and to drive expression of antiviral IFN-stimulated genes (ISGs) [11]. The important roles of IFN-I in host antiviral immunity are explained by the diverse IFN-antagonizing strategies developed in vertebrate viruses. The ability of a given virus to antagonize IFN-I response is an important determination of virus virulence, especially for mosquito and tick-borne arboviruses, such as flaviviruses, as they require viral loads in blood to maintain their vector-host cycles. However, the flaviviruses encompass more than 70 phylogenetically diverse viruses [12], suggesting that the flavivirus-encoded strategies to suppress this critical host response is only beginning to be explored.

Flaviviruses have a single-stranded positive RNA genome, which encodes a polyprotein that is cleaved into three structural proteins of membrane (M), envelope (E), capsid (C), and seven non-structural proteins of NS1, NS2A, NS2B, NS3, NS4A, NS4B, and NS5 [13]. Non-structural proteins play central roles in host innate immune evasion of flaviviruses, in which NS1, NS2B, NS3 and NS5 can suppress IFN-I response via targeting the key molecules of RLR and JAK-STAT signaling. For example, the WNV NS1 interacts with RIG-I and MDA5[14], the DENV NS2B3 cleaves the stimulator of interferon genes (STING) [15]. The YFV, ZIKV, and DENV NS5 inhibits STAT2 through interaction of NS5-STAT2 in a species-dependent manner [16].

The M protein of flaviviruses is synthesized as a precursor membrane (prM) of about 164-168 amino acid (aa) length. The prM protein functions as a chaperon for the folding of E protein, which is cleaved by furin protease just shortly before the virus release [17, 18]. The flavivirus prMs can improve the immunogenicity of E protein and has been used as vaccine component [13, 19]. The TBEV prM (63-69 aa) is important for the association of prM-E protein heterodimers [20], and the 139 and 146 amino acids of ZIKA prM are related to virus replication and pathogenicity in mice [21, 22]. The DENV prM has no effect on IFN-I production [23], and ZIKA prM can inhibit RLR molecules induced interferon production in an unknown manner [24]. Whether flavivirus prMs are involved in host innate immune escape remains to be determined.

Here, we found that TBEV structural proteins prM, C and E can antagonize IFN-I production, in which prM binds to both MDA5 and MAVS, and impedes interaction between MDA5 and MAVS, thereby inhibiting dimerization and nuclear translocation of IRF3. Moreover, flavivirus DENV2 (serotype 2 of DENV), WNV, YFV and ZIKA prM proteins have also been demonstrated to significantly suppress IFN-I production through interacting with MDA5 and/or MAVS. Our findings revealed the immune evasion mechanisms of flavivirus prM proteins, which may contribute to understanding flavivirus pathogenicity, therapeutic intervention and vaccine designation.

## Results

### Flavivirus TBEV viral proteins antagonize IFN-I production

TBEV can suppress host antiviral responses by expressing gene products to inhibit production or signaling of IFNs [25]. We confirmed this phenomenon using the medulloblastoma cell line DAOY, which is susceptible to TBEV infection. The DAOY cells were mock-infected or infected with TBEV at a multiplicity of infection (MOI) of 1.0. In parallel, cells were transfected with the interferon inducer poly (I:C) [26]. The mRNA expression levels of *IFNA* and *IFNB1* were detected upon TBEV infection or poly (I:C) transfection in DAOY cells. Results showed that *IFNA* and *IFNB1* mRNA was gradually increased in a virus dose-dependent manner, whereas it was significantly lower than that of the poly (I:C) transfection group, suggesting that TBEV infection may attenuate host antiviral responses (Supplementary Fig. S1a, b).

To identify the viral proteins of TBEV that inhibit IFNβ production, we constructed the expression plasmids of TBEV proteins and tested their effects on IFNβ promoter activity via luciferase reporter assay upon their expression (Supplementary Fig. S1c). Compared with NS proteins (NS1, NS2A, NS4A, NS4B, and NS5), which have been extensively reported to antagonize IFN-I production in flaviviruses [14, 23, 24, 27], we found the structural proteins prM, C and E could function as interferon antagonists (Fig. 1a). Although the C protein exhibited higher inhibitory effect than prM and E protein, its cytotoxicity was much higher than that of the prM protein (Supplementary Fig. S1d).

**Fig 1.**
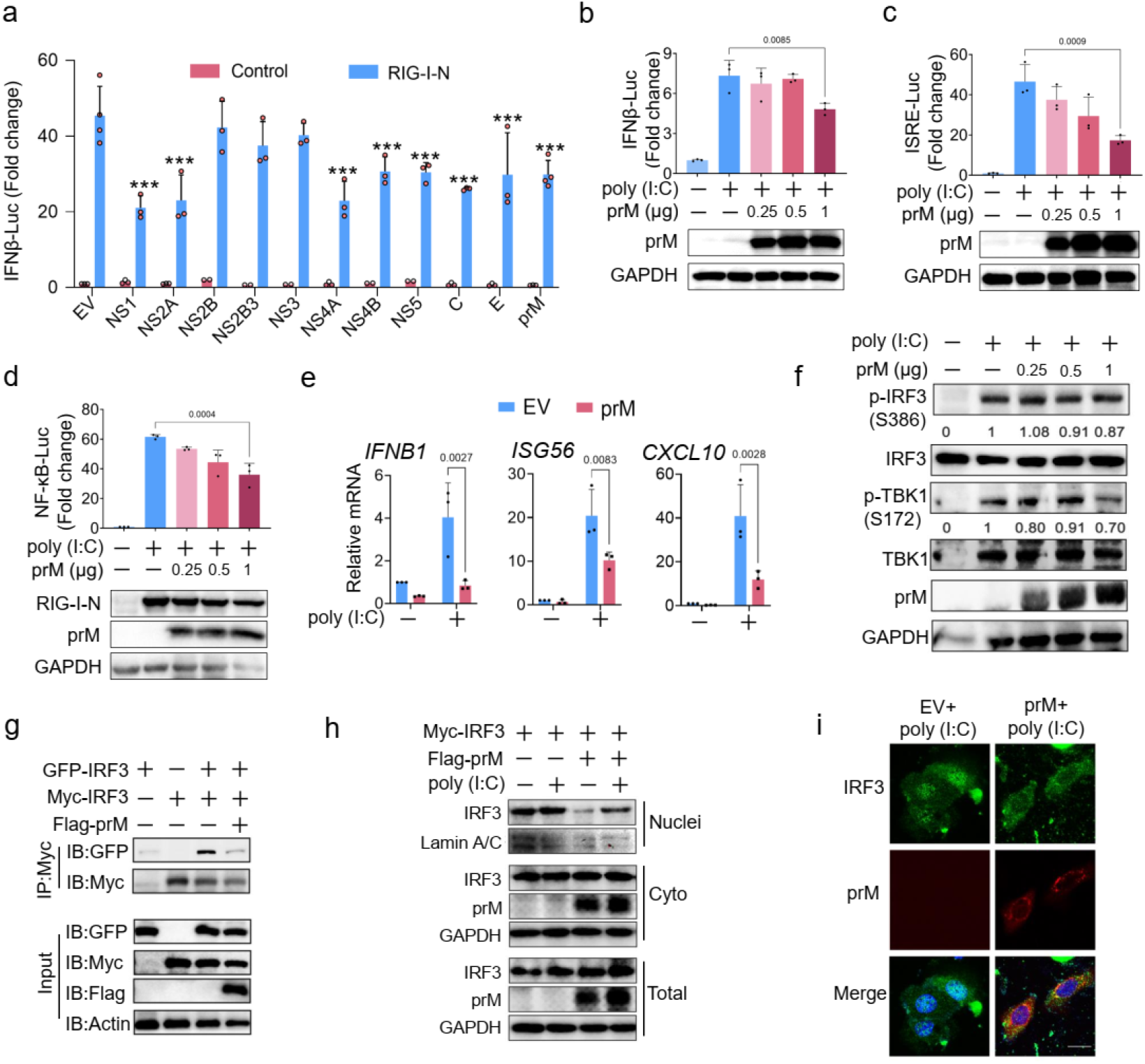
Flavivirus TBEV viral proteins antagonize IFN-I production. (a) HEK293T cells were transfected with empty vector (EV) or plasmids expressing TBEV proteins and RIG-I-N together with IFNβ-Luc and control plasmids, the luciferase activity was measured at 20 hours post-transfection (hpt). (b and c) TBEV prM or EV plasmids, poly(I:C) along with IFNβ-Luc (b) or ISRE-Luc (c) were transfected into HEK293T cells, the luciferase activity was measured at 20 (b) or 24 hpt (c). (d) TBEV prM or EV plasmids, RIG-I-N along with NF-κB-Luc were transfected into HEK293T cells, the luciferase activity was measured at 24 hpt. (e) TBEV prM plasmid along with poly(I:C) were transfected into HEK293T cells, the expression of *IFNB1, ISG56* and *CXCL10* were analyzed by qPCR, GAPDH was used as normalizer. (f) TBEV prM plasmid and poly (I:C) were transfected into HEK293T cells, the cells were harvested at 24 hpt for immunoblot analysis by the indicating antibodies. The relative intensity of phosphorylated IRF3 and TBK1 was calculated using ImageJ software. (g) The Myc-IRF3 and EGFP-IRF3 together with TBEV prM or EV plasmids were co-transfected into HepG2 cells, cells were harvested at 30 hpt, and the cell lysates and immunoprecipitants were analyzed by immunoblot using the indicated antibodies. (h) Myc-IRF3 together with TBEV prM or EV plasmids were co-transfected into HEK293T cells. After 24 h, the cells were activated with poly (I:C) for 8 h. The separated nuclear and cytoplasmic proteins were analyzed for IRF3 by immunoblot. (i) EGFP-IRF3 and TBEV prM plasmids were co-transfected into Hella cells. After 24 h, the cells were activated with poly (I:C) for 8 h and stained with indicated antibodies. Green, IRF3 signal; Red, TBEV prM signal; Blue, DAPI (the nuclear signal). Bar, 10 μm. Bars represent the mean of three biological replicates and all data are expressed as mean ± SE.

The inhibitory effect of TBEV prM was further confirmed by luciferase reporter assay. PrM protein was shown to significantly inhibit the promoter activity of IFNβ and ISRE (IFN-sensitive responsive element) induced by poly (I:C) (Fig. 1b, c), and overexpression of prM suppressed the activity of NF-κB promoter induced by RIG-I-N (the N-terminal CARD domain of RIG-I) (Fig. 1d). TBEV prM also significantly inhibited the mRNA levels of *IFNB1, ISG56*, and *CXCL10* (Fig. 1e).

The activation of IRF3, including its phosphorylation, dimerization and nuclear translocation are important for IFN-I production [28]. We thus examined if TBEV prM suppresses the activation of IRF3. Compared with empty vector, TBEV prM overexpression slightly reduced the phosphorylation of IRF3 and TBK1 induced by poly (I:C) (Fig. 1f), while the dimerization of IRF3 was significantly reduced in the prM transfection group (Fig. 1g). Moreover, the expression of IRF3 in nuclei activated by poly (I:C) was decreased in the prM expression group (Fig. 1h). Consistently, the poly (I:C) induced nuclear translocation of IRF3 was significantly interrupted by the prM expression (Fig. 1i). Taken together, these findings indicated that TBEV prM protein inhibited IFN-I production through inhibiting IRF3 activation.

### Flavivirus TBEV prM protein inhibits RIG-I/MDA5/MAVS induced interferon production

The RLRs members, such as RIG-I and MDA5, are important sensors of cytosolic viral RNA, which play a critical role in IFN-I production (Fig. 2a) [8]. We thus examined the effect of TBEV prM on RLR-mediated IFN-I production, and found that co-expression of TBEV prM suppressed IFNβ promoter activity induced by RIG-I-N, MDA5 and MAVS in a dose-dependent manner (Fig. 2b-d), while overexpression of prM had no significant effect on IFN production induced by TBK1 or IKKε (Fig. 2e, f). Accordingly, the ectopic expression of prM inhibited the mRNA level of *IFNB1*, *ISG56* and *CXCL10* induced by RIG-I-N, MDA5 and MAVS (Fig. 2g-i). As STING has been reported to be involved in signal transduction of RNA virus [29], we also detected the effect of TBEV prM expression on the STING-induced IFN-I production, and found that prM expression had no significant effect on the IFNβ promoter activity or IFNβ and ISGs production induced by STING (Supplementary Fig. S2). These results indicated that TBEV prM inhibited IFN-I production at the MAVS or its upstream level through targeting RIG-I/MDA5/MAVS.

**Fig 2.**
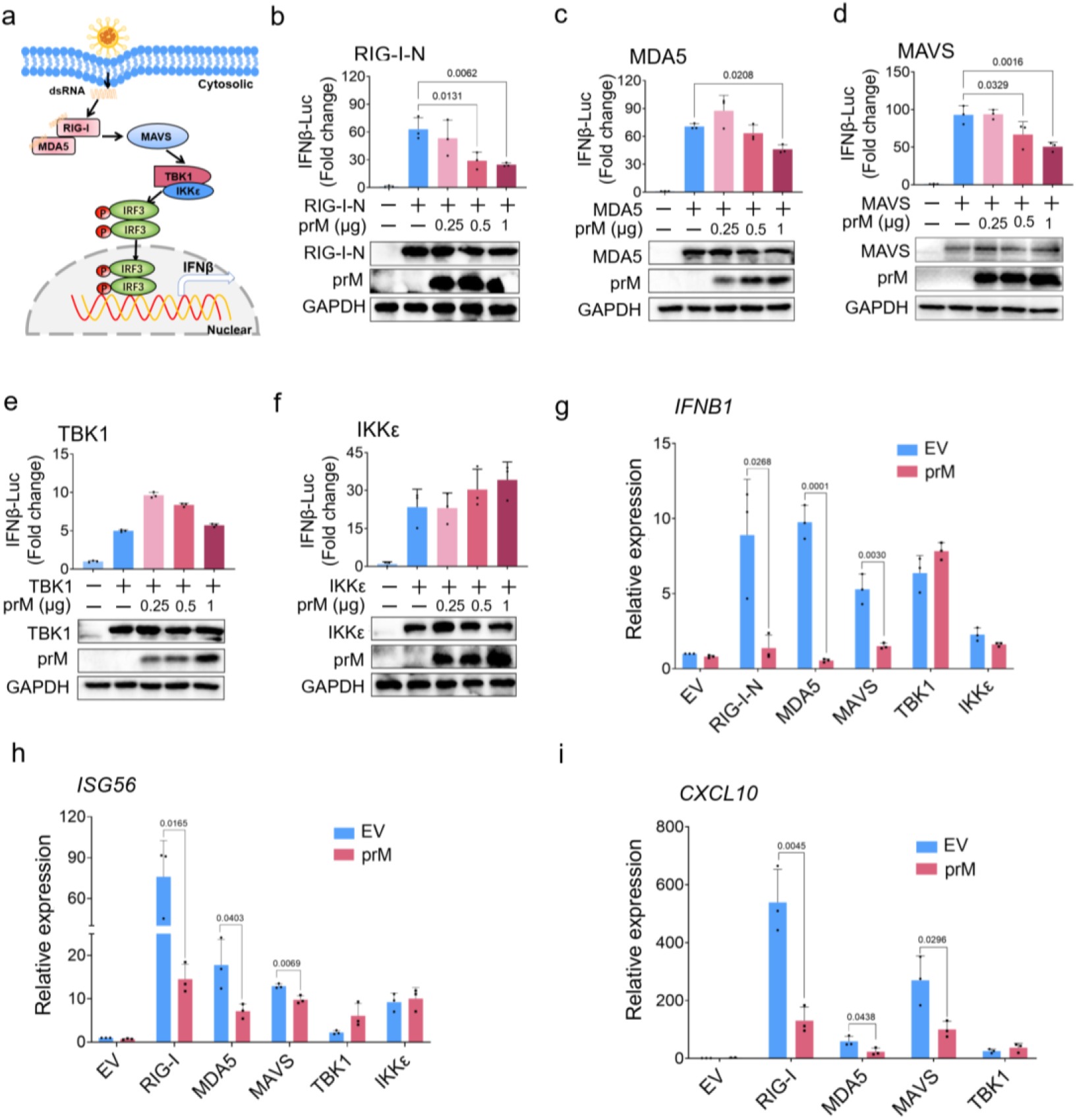
Flavivirus TBEV prM inhibits IFN-I production activated by RIG-I/MDA5-MAVS. (a) Schematic view of RIG-I/MDA5 mediated IFN-I production. (b-f) EV or TBEV prM, the IFNβ-Luc plasmids together with RIG-I-N (b), MDA5 (c), MAVS (d), TBK1 (e) and IKKε (f) were co-transfected into HEK293T cells, the luciferase activity was measured at 20 hpt. The expression of RLRs and prM were detected by immunoblot. (g-i) TBEV prM together with indicated RLR plasmids were co-transfected into HEK293T cells, the expression of *IFNβ* (g)*, ISG56* (h) and *CXCL10* (i) were analyzed by qPCR, GAPDH was used as normalizer. Bars represent the mean of three biological replicates and all data are expressed as mean ± SE.

### Flavivirus TBEV prM interacts with both MDA5 and MAVS

TBEV prM was predicted to have one transmembrane domain on the C-terminal side, and it may localize on the membrane apparatus (Supplementary Fig. S3). We found that TBEV prM mainly localized on endoplasmic reticulum (ER) and partially located on mitochondria apparatus, no obvious localization of prM protein was found on Golgi apparatus (Fig. 3a). As both the ER and mitochondria are important platforms for the signaling transduction of RLRs[30], we examined the co-location of TBEV prM with RLR molecules including RIG-I, MDA5, MAVS, TBK1, and IKKε. Immunofluorescence assay (IFA) results showed that TBEV prM co-localized with RIG-I, MDA5, MAVS and TBK1, but not with IKKε (Supplementary Fig. S4a).

**Fig 3.**
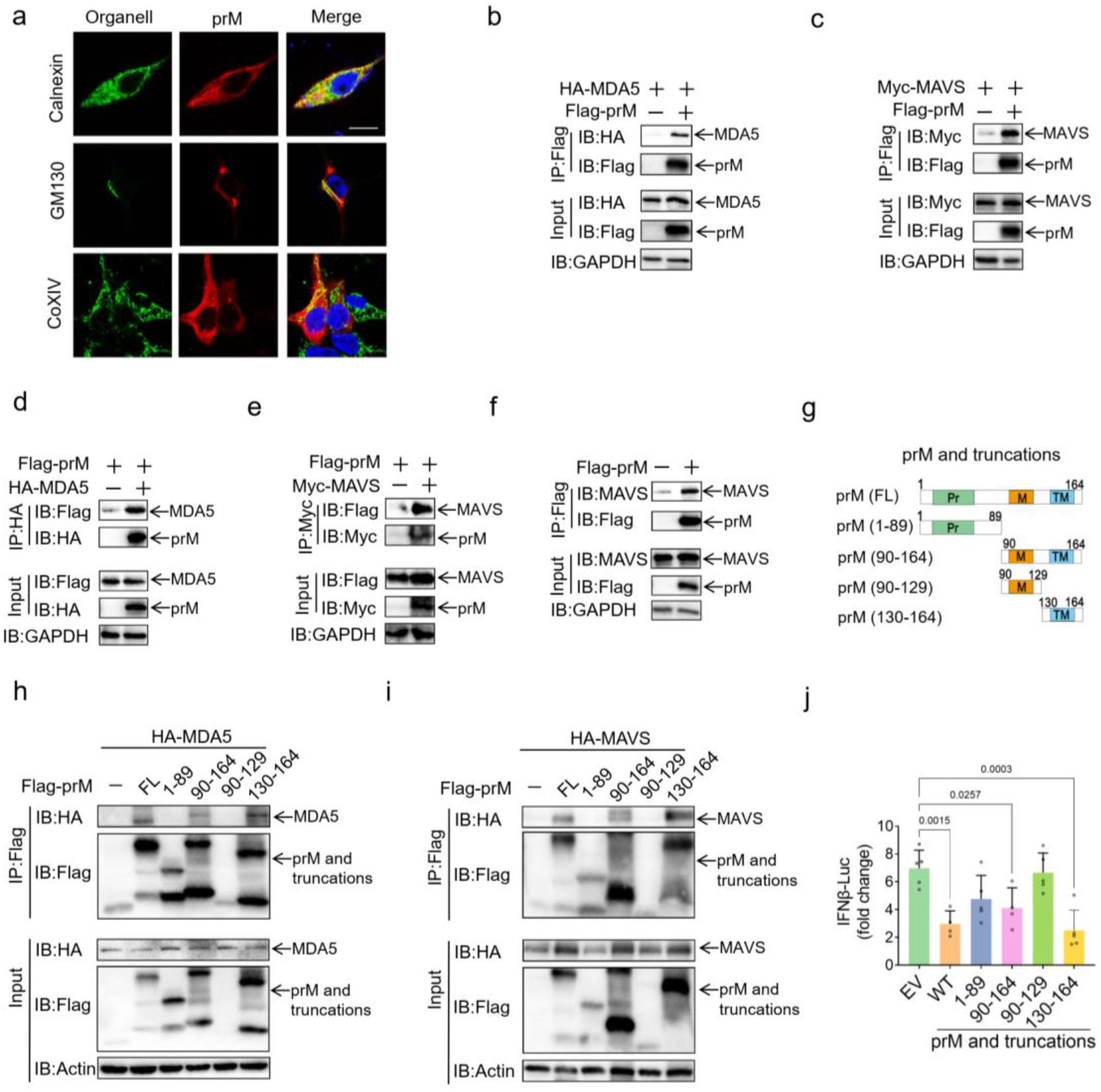
Flavivirus TBEV prM interacts with MDA5 and MAVS. (a) HEK293T cells transfected with TBEV prM plasmid were fixed and stained by Calnexin (endoplasmic reticulum), GM130 (Golgi), COIXV (mitochondria), and Flag antibodies to analyze the location of prM. Green, the corresponding organelles signal; Red, TBEV prM signal; Blue, DAPI (the nuclear signal). Intensity profiles of the indicated proteins were analyzed by Image J line scan analysis. Bar, 10 μm. (b-f) EV or TBEV prM together with HA-MDA5 (b, d), Myc-MAVS (c, e) were transfected into HEK293T cells, cells were harvested at 30 hpt and the cell lysates were co-immunoprecipitated and analyzed by immunoblot using the indicated antibodies. (g) The diagram of TBEV prM truncations. TM, trans-membrane domain. Numbers above the domain names indicate amino acid positions of prM. (h and i) Co-immunoprecipitation and immunoblot analyses of the indicated proteins in HEK293T cells transfected with the full length (FL) and truncated fragments of TBEV prM along with MDA5 (h) or MAVS (i). (j) The expression plasmids of TBEV-prM and its truncations were co-transfected with an IFNβ-Luc and poly (I:C), cells were harvested for luciferase reporter assay at 20 hpt. Bars represent the mean of three biological replicates and all data are expressed as mean ± SE.

Given that TBEV prM can inhibit RIG-I/MDA5-MAVS mediated IFN-I production and co-localized with them, we thus investigated the possible interactions between TBEV prM and RIG-I/MDA5/MAVS. Co-immunoprecipitation showed that upon RIG-I, MDA5 and MAVS co-transfection, both MDA5 and MAVS were found to bind to TBEV prM, which was confirmed by the reverse co-immunoprecipitation (Fig. 3b-e). TBEV prM was also shown to be associated with endogenous MAVS (Fig. 3f), while no obvious interaction was observed between TBEV prM with RIG-I or its downstream TBK1, IKKε, TRAF3, or IRF3 (Supplementary Fig. S4b-f).

We then generated the truncated TBEV prM and determined the domain mapping of TBEV prM with MDA5 and MAVS (Fig. 3g). Results showed that the maturated M (90-164 aa) and TM domain (130-164 aa) of prM could interact with MDA5 and MAVS, while the N-terminal domain (1-89 aa) and the 90-129 aa of prM showed no obvious interaction with MDA5 or MAVS, indicating that the TM domain was mainly responsible for the interaction of prM with MDA5 and MAVS (Fig. 3h, i). Luciferase reporter assay further confirmed that only the maturated M (90-164 aa) and the TM domain (130-164 aa) inhibited poly (I:C) induced IFN-I production (Fig. 3j). Taken together, TBEV prM was shown to interact with both MDA5 and MAVS to antagonize interferon production.

### Flavivirus TBEV prM interferes with the interaction of MDA5 and MAVS

After accepting the signal from RIG-I/MDA5, MAVS aggregates and activates TBK1 and IRF3 to induce interferon production [31]. As TBEV prM interacts with MDA5 and MAVS, we sought to investigate if prM could interfere with the aggregation of MAVS and the complex of MAVS with its up- and down-stream signal molecules. The Flag-tagged TBEV prM, Myc-MAVS together with HA-tagged RIG-I, MDA5, MAVS, TBK1, or TRAF3 expression plasmids were transfected into HEK293T cells, co-immunoprecipitation was conducted using the anti-Myc or HA beads. Results showed that TBEV prM protein impaired the association between MAVS and MDA5 (Fig. 4a). However, prM had no significant effect on the dimerization of MAVS, and did not affect the interaction of MAVS with RIG-I, TBK1, or TRAF3 (Fig. 4b-e). These data indicated that TBEV prM interfered with the recruitment of MAVS by MDA5, consistent with the negative regulation of TBEV prM on the IFN-I production (Fig. 4f).

**Fig 4.**
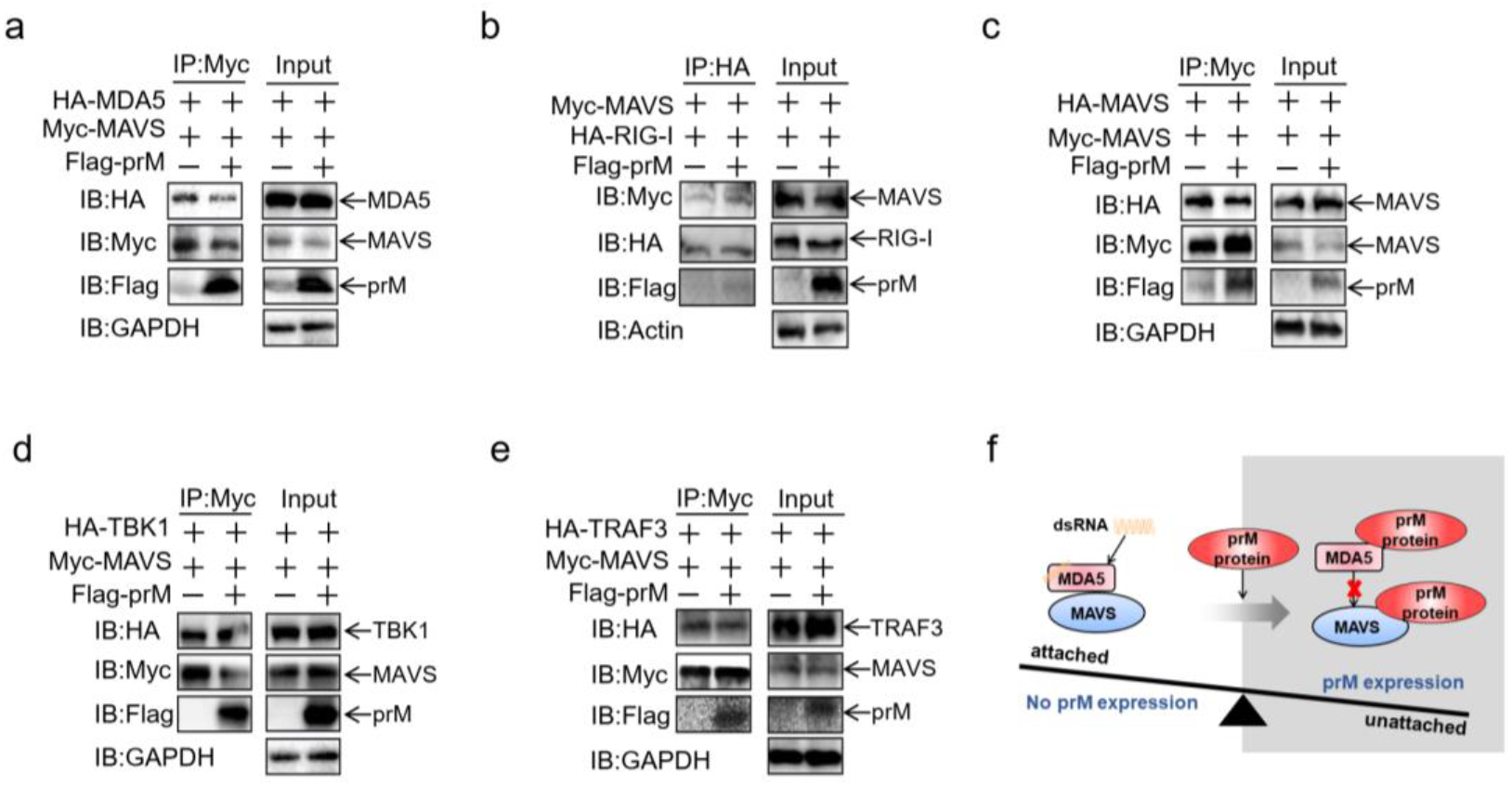
Flavivirus TBEV prM impedes the interaction of MDA5 and MAVS. (a-e) HEK293T cells were transfected with Myc-MAVS together with HA-tagged MDA5 (a), RIG-I (b), MAVS (c) TBK1 (d) or TRAF3 (e) along with EV or Flag-prM. The cell lysates were co-immunoprecipitated and analyzed by immunoblot using the indicated antibodies. (f) Schematic view of TBEV prM interferes with the interaction of MDA5 and MAVS.

### Flaviviruses prM proteins inhibit RIG-I/MDA5-MAVS induced interferon production

With the exception of TBEV, several flaviviruses, including DENV2, JEV, YFV, WNV and ZIKA, also pose severe threats to human health, whose prMs share 14-40% amino acid similarities with that of TBEV (Fig. 5a and Supplementary Table S1). ZIKA prM has shown to suppress IFN-I production, whereas DENV2 prM has no significant effect on interferon production [23, 24]. Next, we investigated if these flavivirus prMs antagonize host innate immunity as the same way as TBEV prM.

**Fig 5.**
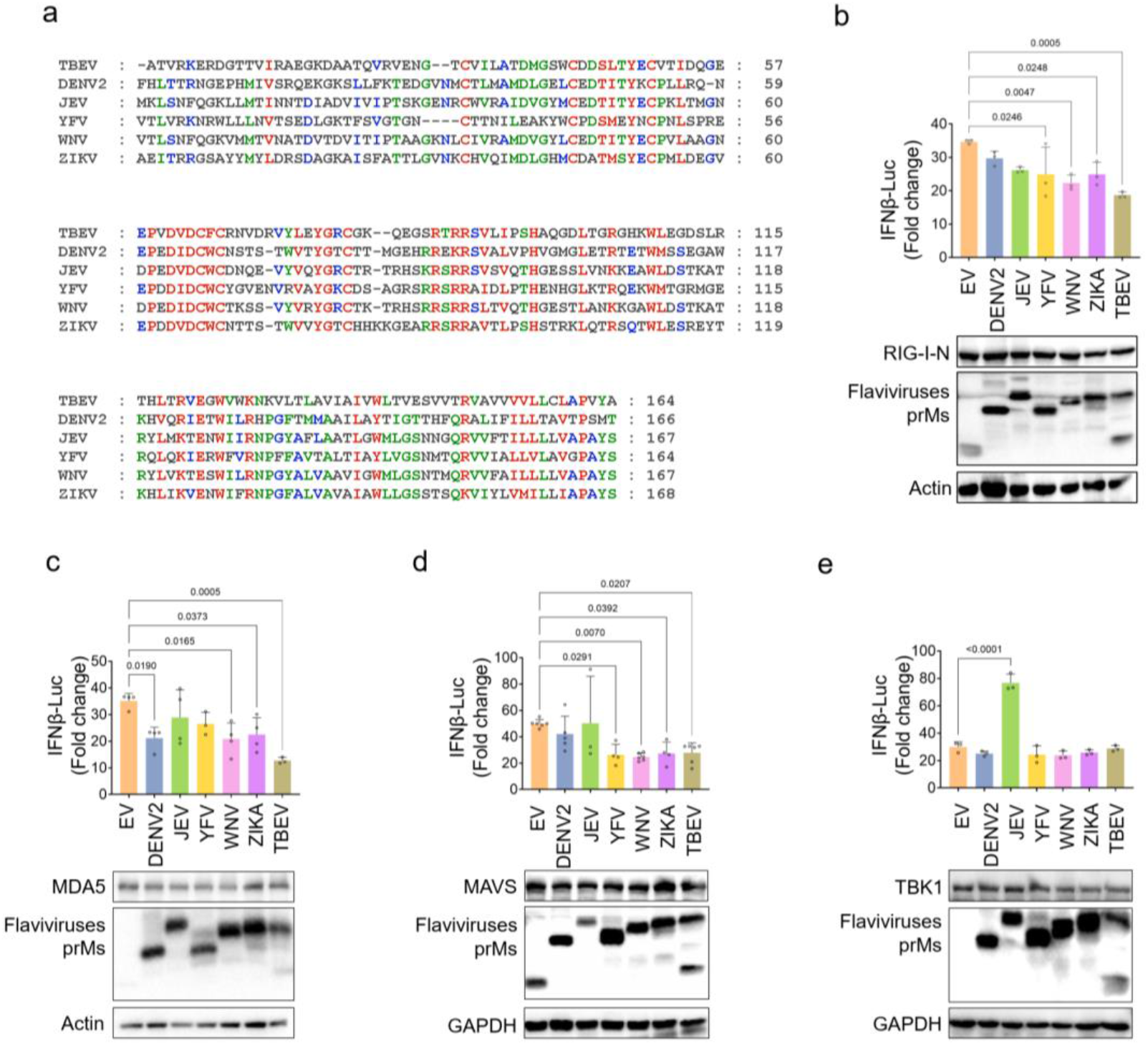
Flavivirus prMs antagonize IFN-I production induced by RIG-I/MDA5-MAVS. (a) The amino acid identity of the prMs from flaviviruses TBEV, DENV2, JEV, YFV, WNV and ZIKA. (b-e) EV or flavivirus prMs, the IFNβ-Luc plasmids together with RIG-I-N (b), MDA5 (c), MAVS (d) and TBK1 (e) were co-transfected into HEK293T cells, the luciferase activity was measured at 20 hpt. The expression of RLRs and prM proteins were detected by immunoblot using the indicated antibodies. Bars represent the mean of three biological replicates and all data are expressed as mean ± SE.

We first cloned the prM of these flaviviruses, and assessed the effect of their expression on IFN-I production induced by RIG-I/MDA5 signaling components. Results showed that the prM protein of YFV, WNV and ZIKA also suppressed the IFNβ promoter activity induced by RIG-I-N, DENV2, WNV and ZIKA prM suppressed the promoter activity of IFNβ induced by MDA5, while prMs of YFV, WNV and ZIKA could inhibit IFN-I induced by MAVS (Fig. 5b-d). None of the flavivirus prMs suppressed the IFNβ production induced by TBK1 (Fig. 5e). Taken together, in contrast with TBEV, the prMs of WNV and ZIKA inhibited IFN-I induced by RIG-I, MDA5 and MAVS, YFV prM protein inhibited IFNβ induced by RIG-I and MAVS, while DENV2 prM only inhibited MDA5 induced IFNβ production, and JEV prM showed no obvious suppression on IFN-I production. These data indicated that prMs of these flaviviruses may inhibit IFN-I production by different mechanisms.

### Flavivirus prM proteins interact with MDA5 and/or MAVS

Given that TBEV prM inhibits IFN-I production via targeting MDA5 and MAVS, we then detected if other flavivirus prMs interact with RLR signaling molecules. The DENV2, JEV, YFV, WNV, ZIKA, and TBEV prM expression plasmids were co-transfected into HEK293T cells with RIG-I, MDA5, MAVS, and TBK1, co-immunoprecipitation showed that flavivirus prMs had no obvious interaction with RIG-I (Fig. 6a), while prMs of DENV2, WNV, ZIKA, and TBEV interacted with MDA5 (Fig. 6d). YFV, WNV, ZIKA, and TBEV prMs could bind to MAVS, with a little stronger interaction for ZIKA and TBEV prMs as comparison with YFV and WNV (Fig. 6c). In contrast, none of the flavivirus prMs interacted with TBK1 (Supplementary Fig. S5).

**Fig 6.**
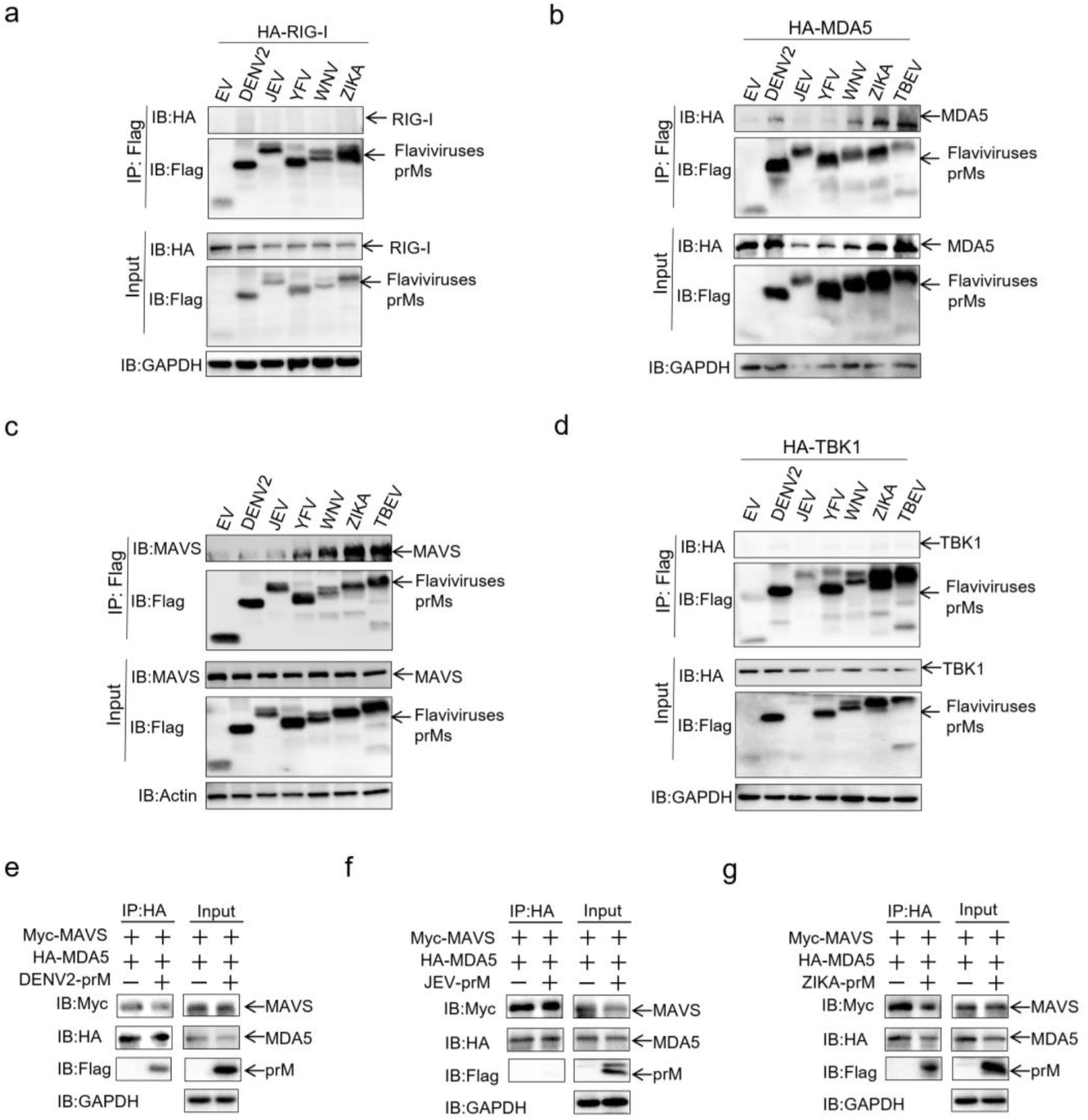
Flavivirus prMs interact with MDA5 and/or MAVS. (a-d) EV or flavivirus prM plasmids together with RIG-I (a), MDA5 (b), MAVS (c) and TBK1 (d) were co-transfected into HEK293T cells. After 30 h, cells were harvested for immunoprecipitant analysis by immunoblot using the indicated antibodies. (e-g) Myc-MAVS, HA-MDA5 together with EV or DENV2-prM (e), JEV-prM (f), and ZIKA-prM (g) were co-transfected into HEK293T cells. After 30 h, cells were harvested for immunoprecipitant analysis by immunoblot using the indicated antibodies.

As TBEV prM interferes with the formation of the MDA5-MAVS complex, we further examined if flavivirus prMs impede the interaction of MDA5 and MAVS. Co-immunoprecipitation showed that WNV and ZIKA prMs that bind to both MDA5 and MAVS significantly affected the formation of the MDA5-MAVS complex (Fig. 6e and Supplementary Fig. S5), while DENV, YFV and JEV prMs that did not interact or only interacted with MDA5 or MAVS did not affect the formation of the MDA5-MAVS complex (Fig. 6f, g and Supplementary Fig. S5). Taken together, TBEV, ZIKA and WNV prMs bind to both MDA5 and MAVS, and interfere with the formation of the MDA5-MAVS complex, while DENV and YFV prMs only interact with MDA5 or MAVS to suppress IFN-I production.

### Flavivirus prM proteins promotes viral replication

We further sought to determine if the flavivirus prM proteins could promote the virus replication. Sendai virus (SeV) is usually used to assess interferon production [32], we hence examined the effects of prM on SeV replication. HEK293T cells transfected with flavivirus prMs were infected with SeV at a MOI of 1.0, and the replication of SeV was determined by IFA, flow cytometry, or immunoblot assay. IFA showed that TBEV prM protein significantly promoted the replication of SeV, higher in 1 μg than in 0.5 μg prM transfection group (Fig. 7a). Flow cytometry analysis showed that along with the increasing dose of prM transfected, the percentage of SeV-positive cells was gradually elevated, significantly higher in the 1.0 μg prM transfection group as comparison with the control group (Supplementary Fig. S6). Immunoblot analysis also showed a significantly higher SeV protein levels in prM transfection group compared with the empty vector and control group (Fig. 7b). The TM and mature M domain that interacted with MDA5 and MAVS obviously promoted SeV replication (Fig. 7c). Given the interferon antagonizing activity of the flaviviruses, we further detected the effect of prMs from DENV2, JEV and ZIKA on SeV replication. Similar to TBEV, ZIKA prM significantly promoted replication of SeV, while DENV2 and JEV prM proteins showed no significant effect (Fig. 7d). Consistently, flow cytometry analysis revealed that the full length and the 130-164 aa truncation of TBEV and ZIKA prMs significantly enhanced the percentage of SeV-positive cells, while the SeV-positive cells in DENV2 prM transfection group was a little bit higher than the empty vector group, and JEV prM did not affect SeV production (Fig. 7e). Taken together, TBEV and ZIKA prMs can facilitate SeV replication.

**Fig 7.**
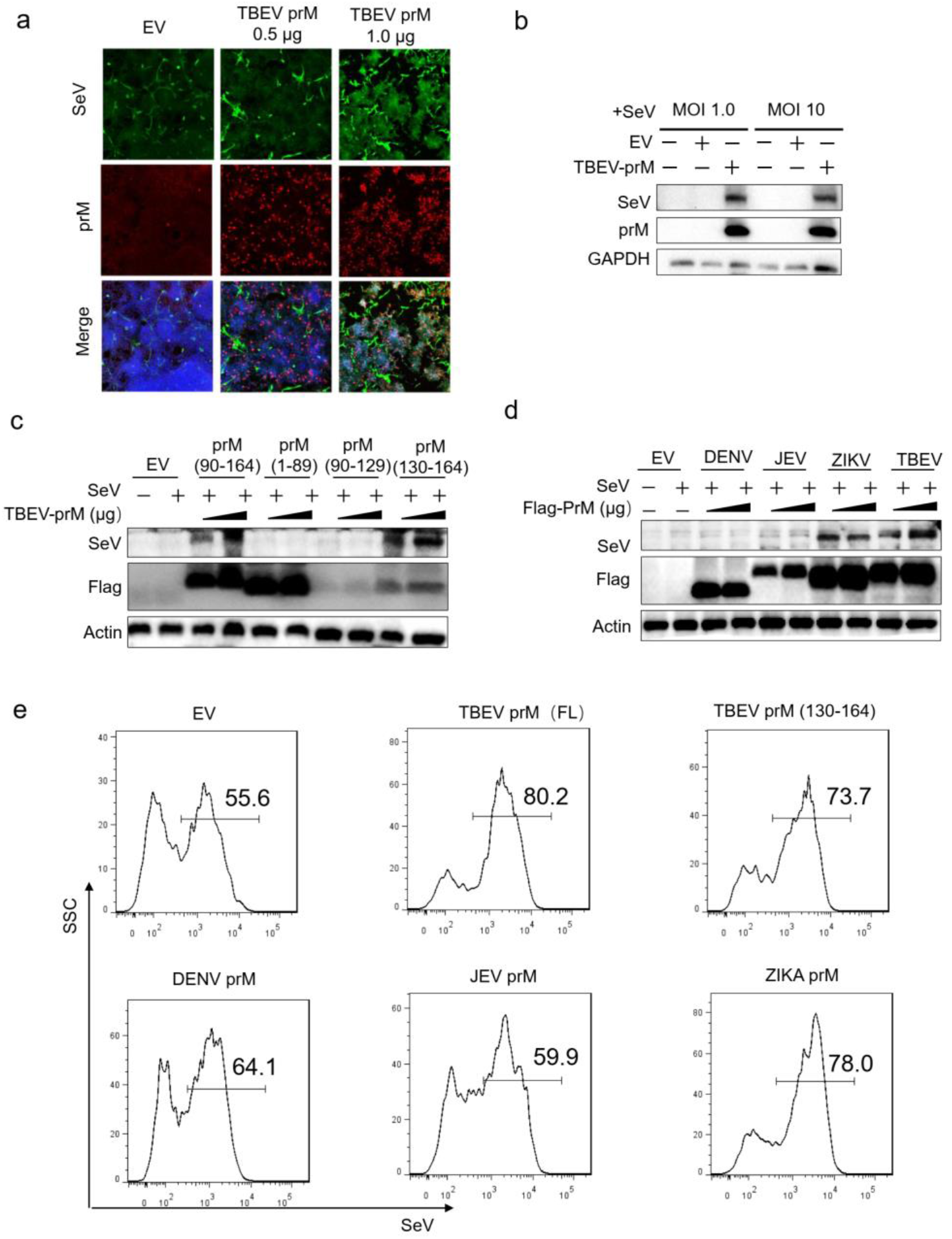
Flavivirus prM proteins facilitate SeV replication. (a) EV or TBEV prM plasmids were transfected into HEK293T cells. After 24 h, cells were infected with SeV (MOI 1.0) for another 20 h, the replication of SeV was analyzed by IFA analysis with anti-SeV and -Flag antibodies. (b-e) EV and full length (FL) of TBEV prM (b-e), TBEV truncation (c, e) or flavivirus prM (d, e) plasmids were transfected into HEK293T cells. After 24 h, cells were infected with SeV (MOI 1.0 and 10). The cells were harvested after 20 h and analyzed by immunoblot (b-d) and flow cytometry analysis (e).

## Discussion

All vector-borne flaviviruses studied till now need to overcome the antiviral innate immunity, particularly IFN-I responses, to infect vertebrate host. The non-structural proteins of flaviviruses are mainly involved in viral replication and host innate immune escape, and the structural proteins are responsible for the virus assembly. In this study, we found that TBEV structural proteins prM, C and E could antagonize IFN-I production, in which prM interacted with both MDA5 and MAVS to inhibit RLR antiviral signaling. Interestingly, ZIKA and WNV prMs were also demonstrated to interact with both MDA5 and MAVS, while dengue virus serotype 2 (DENV2) and YFV prMs associated only with MDA5 or MAVS to suppress IFN-I production. In contrast, JEV prM could not suppress IFN-I production. These findings of immune evasion mechanisms mediated by prM of flaviviruses help to explain the pathogenicity of emerging flaviviruses.

Several innate immune escape strategies have been identified in TBEV, which are associated with the delayed interferon response during the infection. TBEV prM, NS1, NS2A, and NS4B can substantially block the transcriptional activity of IFN-β promoter [25]; NS4A binds STAT1 and STAT2 to suppress their phosphorylation and dimerization, thereafter inhibiting IFN-I signaling [33]; NS5 associates with membrane protein scribble to impair interferon-stimulated JAK-STAT signal [34]; NS5 activates IRF3 in a manner dependent on RIG-I/MDA5 [35]. TBEV NS5 interacts with the prolidase, a host protein required for maturation of IFNAR1 [36], NS5 has also been shown to interact with the mammalian membrane protein Scribble that is implicated in T cell activation [37]. Our findings further support that TBEV non-structural proteins NS1, NS2A, NS4A, NS4B, and NS5 act as interferon antagonists during TBEV infection. Importantly, we found that TBEV structural proteins, including prM, C, and E, play an important role in avoiding host innate immunity, and prM interact with both MDA5 and MAVS to inhibit IFN-I production. However, anti-interferon mechanisms of TBEV C and E proteins remain to be further investigated.

Our study also demonstrated that the prM proteins of WNV and ZIKA inhibited IFN-I induced by RIG-I, MDA5 and MAVS, prM of YFV inhibited IFNβ induced by RIG-I and MAVS, while DENV2 prM only inhibited MDA5-induced IFNβ production, and JEV prM showed no obvious suppression on IFN-I production. Mechanically, WNV and ZIKA prMs interact with both MDA5 and MAVS, whereas DENV2 and YFV prMs bind to MDA5 or MAVS to evade innate immunity.

The prM protein of flavivirus is involved in the virus life cycle, whose N-glycosylation is essential for protein trafficking and folding and virion assembly [38–40]. DENV2 prM plays a crucial role in the viral assembly [41]. ZIKA prM is associated with the virus growth and pathogenesis in mice [21], and the single S139N mutation in the prM is associated with fetal microcephaly [22]. Flavivirus virulence and replication efficiency positively correlate with the ability to inhibit the IFN-I production and signal. TBEV virulence is associated with the ability to suppress IFN-I production [25]. Therefore, our findings of prM proteins to subvert host innate immunity would contribute to further clarification of the pathogenesis of flaviviruses. In fact, viral structural proteins suppress the IFN-I antiviral response, such as SARS coronaviruses, Foot-and-mouth disease virus, Bluetongue virus, and Ebola virus [26,32,42–44].

Previous studies have demonstrated that a prM-E DNA vaccine can offer complete protection against ZIKV challenge, whereas a prM-deleted mutant plasmid DNA vaccine cannot provide the same protection, suggesting that prM can effectively enhance vaccine immunogenicity of flaviviruses [45]. A better understanding of the role of prM protein in pathogenesis of flaviviruses in this study has important implications in the development of new antiviral drugs and vaccines. In summary, our findings revealed that flavivirus prM proteins inhibit IFN-I production via interacting MDA5 and/or MAVS in a species-dependent manner, which may contribute to understanding flavivirus pathogenicity, therapeutic intervention and vaccine development.

## Materials and methods

### Cells and viruses

Human embryonic kidney HEK293T cells, hepatic carcinoma HepG2 cells, Hella cells, Vero cells, and human medulloblastoma tumor DAOY cells were cultured in DMEM (HyClone, Logan, USA) supplemented with 10% FBS (BBI, Shanghai, China) and 1% penicillin-streptomycin (100 IU/ml) at 37°C with 5% CO_2_. All cells were tested negative for mycoplasma.

TBEV, belonging to the Far Eastern (FE) subtype, was isolated from *Ixodes persulcatus* ticks in northeast China[46] and Sendai virus (SeV) was kindly provided by Professor Siyang Huang at Yangzhou University. TBEV and SeV were propagated in Vero cells.

### Plasmid construction and antibodies

TBEV structural and nonstructural proteins (GenBank access number MN615726.1), DENV2 (NC_001474), JEV (NC_001437.1), YFV(NC_002031.1), WNV(NC_001563.2), ZIKA (NC_012532.1) prM genes or TBEV prM truncations were cloned into Flag-VR1012 at Sangon Biotech (Shanghai, China). The expression plasmids of IFN-β-Luc reporter, ISRE-Luc reporter, Renilla-Luc, RIG-I, MDA5, TBK1, IKKε, IRF3 and TRAF3 were constructed in our previous study [47]. The NF-κB-Luc and MAVS plasmids were purchased from Miaoling Biotech (Wuhan, China). Anti-HA, anti-GST, anti-Flag, anti-Myc, anti-GAPDH, anti-Actin CoraLite 594-conjugated IgG, and CoraLite 488-conjugated IgG secondary antibodies were obtained from Proteintech (Wuhan, China); anti-Lamin A/C antibody from TransGen (Beijing, China); anti-Phospho-IRF3 (S396) and anti-Phospho-TBK1 antibodies from Cell Signaling technology (Danvers, USA); and anti-SeV antibody from MBL (Beijing, China).

### Luciferase reporter assay

The dual-Luciferase reporter assay system (Promega, Madison, USA) was used for luciferase assays. HEK293T cells were seeded in 24-well plates (2×10^5^ cells/well) and transfected with luciferase reporter and control Renilla-Luc plasmids combined with target plasmids, the luciferase activity of IFNβ-Luc and ISRE-Luc was detected at 20 or 24h post-infection.

### Cell viability assay

HEK293T cells were seeded 7000 cells/well in opaque-walled white 96-well plate. After incubation for 24 h, 150 ng control plasmid and one of the TBEV gene expression plasmids was transfected into the cells. Following incubation for another 48 h at 37 °C, cell viability was determined using a CellTiter-Glo 2.0 Assay (Promega, Madison, USA).

### Coimmunoprecipitations and immunoblot assay

After 30 h post-transfection, cells were lysed in a immunoprecipitation (IP) lysis buffer containing 50 mM Tris (pH 7.5), 1 mM EGTA, 1 mM EDTA, 1% Triton X-100, 150 mM NaCl, 100 μM phenylmethylsulfonyl fluoride (PMSF) and complete TM protease inhibitors (Selleck, Houston, USA) for 30 min at 4°C. Cell lysates were incubated overnight with ANTI-FLAG^®^ M2 Affinity Gel (Sigma-Aldrich, St. Louis, USA), EZview™ Red Anti-HA Affinity Gel (Millipore, Billerica, USA), or Anti-MYC Affinity Gel (Millipore, Billerica, USA), then proteins were separated by SDS-PAGE and transferred onto PVDF membranes. After blocking in TBST containing 5% BSA, the blots were probed with primary antibodies, and the relative band intensities were determined with ChemiDoc XRS+ Molecular Imager software (Bio-Rad, Philadelphia, USA).

### Quantitative PCR (qPCR)

Total cellular RNA was isolated with EasyPure^®^ RNA Kit (TransGen, Beijing, China). Viral RNA in culture supernatants was extracted using TIANamp Virus RNA Kit (Tiangen, Beijing, China), and the first-strand cDNA was synthesized by Trans Script First-Strand cDNA Synthesis Super Mix (TransGen, Beijing, China). The qPCR was conducted with SYBR Green Master (Roche, Rotkreuz, Switzerland) as previously described [48]. The results were normalized by the house-keeping gene GAPDH.

### Immunofluorescence assay (IFA)

HEK293T or HepG2 cells cultured on 12-mm coverslips were transfected with indicated plasmids. After 24 h, cells were fixed with 4% paraformaldehyde, and permeated with 0.5% Triton X-100. The cells were washed with PBST, blocked in 1% BSA, and stained with primary antibodies, followed by staining with CoraLite 594 or CoraLite 488 conjugated IgG secondary antibodies. Nuclei were stained with DAPI (Yesen Biotechnology, Shanghai, China). Fluorescence images were obtained and analyzed using a confocal microscope (FV3000, OLYMPUS).

### Nuclear and cytoplasmic extraction

The nuclear and cytoplasmic fraction of indicated cells were separated using the nuclear and cytoplasmic protein extraction kit (Beyotime, China) according to the manufacturer’s instructions. The purified cytoplasmic and nuclear fraction were subjected to immunoblot with the relevant antibodies.

### Flow cytometry

HEK293T cells were harvested, and washed with PBS. After staining with an anti-SeV antibody, cells were analyzed using a BD LSRFortessa flow cytometer. Data analysis was carried out with the FlowJo software.

### Statistical analysis

The results are representative of at least three independent experiments and shown as the mean ± SD values. For statistical analysis, two-tailed unpaired Student’s t tests were performed in GraphPad Prism 9.0.2, and P value of less than 0.05 was considered statistically significant.

## Data availability

All data supporting the findings of this study are available within the article and its supplementary information or from the corresponding author upon reasonable request.

## Supporting information

**S1Table. The prM amino acid similarities of several flaviviruses.**

**S1 Fig. Flavivirus TBEV proteins inhibit interferon production.** (a and b) DAOY cells were mock-infected or infected with TBEV (JL-T75) at a MOI of 1.0, 2.0 and 4.0 or activated by Poly (I:C) for 12 h, cells were collected and the mRNA level of *IFNα* and *IFNβ* were detected by qPCR, GAPDH were using as control. (c) The immunoblot analysis of the 11 TBEV proteins. Red boxes indicated the viral proteins of TBEV. (d) Cellular toxicity of TBEV proteins. HEK293T cells were transfected with 150 ng of plasmid for each TBEV expression plasmids, cell viability was analyzed by a luminescent cell viability assay. Cell viability <70% were indicated for red color.

**S2 Fig. Flavivirus TBEV prM had no significant effect on STING induced IFN-I production.**

(a) TBEV prM or EV, the IFNβ-Luc plasmids together with STING were co-transfected into HEK293T cells, the luciferase activity was measured 20 hpt. The expression of STING and prM were detected by immunoblot assay. (b) 0.5 μg of prM and STING plasmids were co-transfected into HEK293T cells, the expression of *IFNβ, ISG56* and *CXCL10* were analyzed by qPCR, GAPDH was used as normalizer. Bars represent the mean of three biological replicates and all data are expressed as mean ± SE.

**S3 Fig. The TBEV prM protein is predicted to contain one transmembrane motif.** (a) The transmembrane motifs in the TBEV prM protein were predicted by the TMHMM server, version 2.0. (b) The transmembrane motif is from 130 to 152aa in the TBEV prM protein.

**S4 Fig. Flavivirus TBEV prM does not interact with RIG-I/TBK1/IKKε/TRAF3/IRF3.** (a) Flag-prM and RLRs expression plasmids were transfected into HEK293T cells. After 24 h, the cells were fixed and stained by Flag and HA antibodies to analyze the co-location of prM and RLRs. Green, RLR proteins signal; Red, TBEV prM signal; Blue, DAPI (the nuclear signal). Intensity profiles of the indicated proteins were analyzed by Image J line scan analysis. (b-f) HEK293T cells were transfected with Flag-prM or EV together with HA-RIG-I (b), HA-TBK1 (c), HA-IKKε (d), HA-TRAF3 (e) or Myc-IRF3 (f), cells were harvested 30 hpt and the cell lysates were co-immunoprecipitated and analyzed with the indicated antibodies.

**S5 Fig. WNV prM interferes with the complex of MDA5 and MAVS.** Myc-MAVS, HA-MDA5 together with EV or WNV-prM (a) and YFV-prM (b) were co-transfected into HEK293T cells. After 30 h, cells were harvested and the cell lysates were co-immunoprecipitated with anti-HA antibody. The cell lysates and immunoprecipitants were analyzed by immunoblot using indicated antibodies.

**S6 Fig. The TBEV prM protein facilitate SeV replication.** (a) EV and TBEV prM plasmids were transfected into HEK293T cells, cells were infected with SeV (MOI 1.0) after 24 h, the replication of SeV was analyzed by flow cytometry analysis. (b) Two independent experiments were conducted in A and the data were showed in column graph. *p < 0.05, **p < 0.01.

## Acknowledgments

This work was supported by grant from National Natural Science Foundation of China (81972873 and 82002165) and the Pearl River Talent Plan in Guangdong Province of China (2019CX01N111).

## Author Contributions

**Conceptualization:** Quan Liu, Zedong Wang.

**Funding acquisition:** Quan Liu.

**Investigation:** Liyan Sui, Yinghua Zhao, Wenfang Wang, Hongmiao Chi, Tian Tian, Ping Wu, Jinlong Zhang, Yicheng Zhao, Zheng-Kai Wei, Zhijun Hou, Guoqiang Zhou, Guoqing Wang.

**Methodology:** Liyan Sui, Yinghua Zhao.

**Project administration:** Quan Liu.

**Writing – original draft:** Liyan Sui, Yicheng Zhao.

**Writing – review & editing:** Quan Liu, Yicheng Zhao.

